# Robust Perfect Adaptation in Biomolecular Reaction Networks

**DOI:** 10.1101/299057

**Authors:** Fangzhou Xiao, John C. Doyle

## Abstract

For control in biomolecular systems, the most basic objective of maintaining a small error in a target variable, say the expression level of some protein, is often difficult due to the presence of both large uncertainty of every type and intrinsic limitations on the controller’s implementation. This paper explores the limits of biochemically plausible controller design for the problem of robust perfect adaptation (RPA), biologists’ term for robust steady state tracking. It is well-known that for a large class of nonlinear systems, a system has RPA iff it has integral feedback control (IFC), which has been used extensively in real control systems to achieve RPA. However, we show that due to intrinsic physical limitations on the dynamics of chemical reaction networks (CRNs), cells cannot implement IFC directly in the concentration of a chemical species. This contrasts with electronic implementations, particularly digital, where it is trivial to implement IFC directly in a single state. Therefore, biomolecular systems have to achieve RPA by encoding the integral control variable into the network architecture of a CRN. We describe a general framework to implement RPA in CRNs and show that well-known network motifs that achieve RPA, such as (negative) integral feedback (IFB) and incoherent feedforward (IFF), are examples of such implementations. We also develop methods to solve the problem of designing integral feedback variables for unknown plants. This standard control notion is surprisingly nontrivial and relatively unstudied in biomolecular control. The methods developed here connect different existing fields and approaches on the problem of biomolecular control, and hold promise for systematic chemical reaction controller synthesis as well as analysis.

## I. INTRODUCTION

Control theory and engineered genetic circuits have been used to control behavior of cells or cell populations [1], [2], [3], [4]. Since disturbances, uncertain dynamics, and stochastics are particularly severe in biomolecular systems [5], the simple notion of perfect adaptation has been a focus in the biological control community [6], [7].

Perfect adaptation (PA) describes the property that the output or error of the system goes to zero for a set of initial conditions and constant disturbances. Robust perfect adaptation (RPA) then describes the property that the system has PA despite uncertain parameters or uncertain dynamics. PA is the same as constant disturbance rejection known in control theory community, and it has long been studied in traditional control disciplines such as electrical and mechanical engineering [8]. The internal model principle in control theory characterizes that a system has PA if and only if there exists an integral control variable in the system [9], [10]. If integral control is a structural property of a system, not dependent on uncertainty, then it also implies RPA, almost trivially. This is easily implemented in electronics, but as we will see, not in biochemical controllers.

RPA is central to biological systems, from chemotaxis[11], [12] to glycolysis [13], from maintenance of homeostasis [14] to cell signaling [15]. In pursuit of the principles behind PA in biological systems, previous studies have focused on discovering biomolecular network topologies that could achieve perfect adaptation through extensive computational studies or case studies in specific biological systems [16], [17].

Dual to the systems biology pursuit in discovering principles underlying PA, synthetic biologists have focused on designing robust integral control variables that are implementable by biomolecular reactions to achieve RPA in engineered cells [6]. RPA is a particularly attractive property for synthetic biologists as it is difficult in general to have complete knowledge of the plant or to construct precise controller dynamics. Because detailed and precise models are rare in biology, these integral variable design problems are usually posed for unknown plants. The constraint that the integral variable is implementable by biomolecular reactions is essential, as it then can be implemented by inserting genetic circuits into cells to achieve PA for variables of interest.

Most previous studies could be considered “reverse engineering”, where examples and case studies of natural biological systems are thoroughly investigated to accumulate heuristics and principles. Here we take a more “forward engineering” approach. We use chemical reaction networks (CRNs) as the general model for what biomolecular systems are capable of implementing [18], [19], [20]. Starting from physical constraints on the types of dynamics that CRNs can take, we describe a general framework to find implementations of (R)PA in CRNs. Focusing on stochastics, [21] employs a similar constraint-focused approach to find design principles for noise suppression [22].

The theoretical tools used in this paper are rather elementary. However, we are able to describe perfect adaptation in biomolecular reaction networks through a constraint-focused control theory framework that builds on and connects different existing fields and approaches. This work also tries to emphasize the different focuses and questions asked in biomolecular control compared to the more common perspectives in control theory for engineering systems.

We first briefly review the most relevant control theory results on perfect adaptation in Section II, and show how (R)PA is obtained in simple electrical circuits. In Section III-A, we show that, due to physical constraints on CRN dynamics, integral control variables cannot be easily implemented using the dynamics of chemical species. This forces biological systems to encode integral variables in more complex biomolecular network architecture. In Section III-C, we develop three general approaches to implement (R)PA in CRNs, with previously reported motifs as examples, connecting results from systems biology with results from chemical reaction network theory. In Section III-E, we address the question of designing integral variables for unknown plants, where sequestration feedback circuit [6] is one example. In Section III-F, we discuss how designing a incoherent feedforward architecture fundamentally requires knowledge about the plant and how our framework can be used to derive such architectures.

## II. ROBUST PERFECT ADAPTATION

The notion of perfect adaptation (PA) comes from systems biology [11]. It is equivalent to constant disturbance rejection in control theory, and has been well studied [9], [10], [12]. For completion, we define PA for the type of systems studied here, and review related key results and relate it to incoherent feedforward (IFF).

Consider a deterministic control-affine SISO closed loop system with state 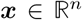, disturbance 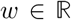, and output 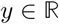, with dynamics

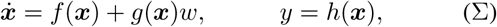

where 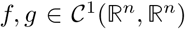, and 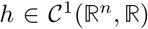 are functions with continuous derivatives.

### Definition 1

System (Σ) is said to satisfy perfect adaptation (PA) with open subsets 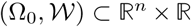, if *y(t)* → 0 as *t* → ∞ for all initial conditions *x*(0) ∈ Ω_0_ and constant disturbances 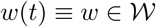.

We often want PA in spite of parameter variations or uncertain dynamics. This corresponds to robust perfect adaptation (RPA).

### Definition 2

System (Σ) is said to satisfy robust perfect adaptation (RPA) with open subsets 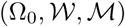 where 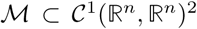, if the system is PA with 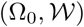 for all 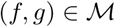.

The key result in control theory on perfect adaptation is the internal model principle. For constant disturbance rejection, the internal model principle states that the system (Σ) needs to satisfy integral feedback. The system (Σ) is said to have integral feedback if there exists a function of *x* such that it is equal to time integral of *y*. More formally,

### Definition 3

System (Σ) is said to have integral feedback (IFB) if 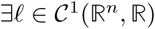 such that z = *ℓ(x)* satisfy 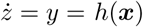.

### A. PA in linear systems

PA in linear systems was studied in the classic work [9]. We focus on the case of SISO systems. To build intuition, we show how PA is equivalent to integral feedback, and how robustness comes naturally as a consequence of stability.

Consider a SISO LTI system,

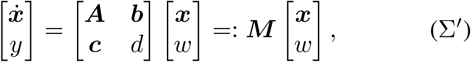

where 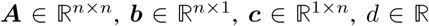. Since the system is linear, the set of initial conditions and disturbances in the definition of PA can always be 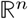 and 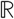. The definition of RPA then becomes

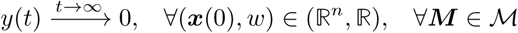

for some open subset of parameters 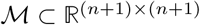.

Assuming the system is stable, i.e. *A* is Hurwitz, then it reaches steady state. At steady state, we have *Ax* + *bw* = 0, so *x* = −*A^−1^bw.* Therefore *y* = (d − *cA^−1^b)w* for any constant *w*, which implies *d − cA^−1^b* = 0. This characterization has an immediate incoherent feedforward (IFF) interpretation. When *d* ≠ 0, disturbance *w* enters output *y* through both *d* and *cA^−1^b* terms, and they cancel each other in steady state. This type of architecture, termed IFF in systems biology literature, has been widely found in biological regulatory systems and is considered an equally important architecture to IFB [23]. Note, however, that if *d* ≠ 0, an IFF interpretation is not as clear as IFB.

By the Schur complement formula for determinant, we see that

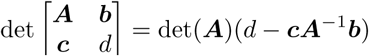

which is 0 if and only if *d* − *cA^−1^b* = 0. So, assuming system (Σ′) is stable,

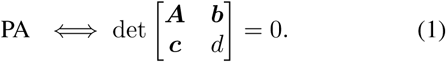

Note that checking whether a set of matrices has zero determinant has efficient probabilistic algorithms [24], so for linear systems we can efficiently check whether a set of systems 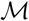 satisfy RPA.

Now we show that PA is equivalent to integral feedback (IFB). Consider ***k* = *cA^−1^.*** Then *z* = ***kx* = *cA^−1^x*** satisfy 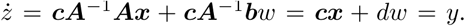. So, assuming the system (Σ′) is stable,

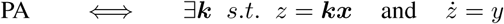

Here the existence of this *z* variable shows the system (Σ′) satisfies IFB. This is a special case of the internal model principle [9], as *z* is a model of all possible constant inputs when *y* = 0. Note that, in comparison to IFF, the IFB interpretation is not as natural, as it requires a change of coordinates, but the IFB argument holds without change when *d* = 0.

Now we consider robustness. From (1), we see that if ***A*** is Hurwitz and *M* satisfy det *M* = 0, then PA is true. Therefore, system (Σ′) is RPA wilh set 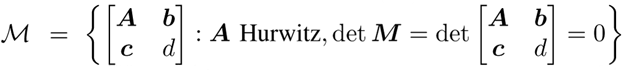.

We might consider the zero determinant condition too restrictive. However, when we consider ***M*** matrices that arise from physical systems, the zero determinant condition can easily be encoded in the physical interconnections of the system, so that any physical parameter variations would preserve the zero determinant condition. Therefore, as long as parameter uncertainties preserve stability of A, the PA property of system (Σ^′^) is robust to such uncertainty. To better illustrate this point, we consider RPA in a simple RLC electrical circuit.

### B. RPA in electrical systems

Assume the RLC circuit in Figure 1 has output *v*, the (unique) voltage of the circuit, and input *i*_*s*_, the source current. The system can be written in the form of (Σ^′^) is

**Fig. 1.**
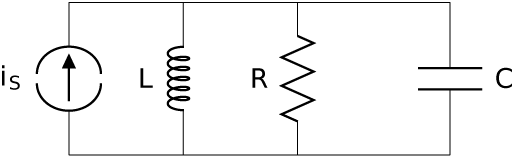
Schematic diagram showing a simple RLC circuit. The input is a source current *i*_*s*_, and output is voltage across the circuit components.

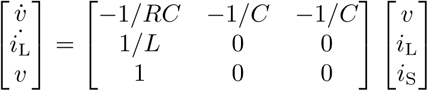

So det *M =* 0 with *i*_*L*_ as the integral variable of output *v*. We first notice that integral variables are easy to construct in electrical circuits because natural integrator elements such as capacitors and inductors exist. This is also the case for cyberphysical systems we commonly face nowadays, with integration easily performed by embedded computers on any observed signals. We will see in Section III-A, however, that this is not the case for biomolecular systems.

We also comment that the sparsity pattern of the *M* matrix in this example is *physical*. This means the zeros in the *M* matrix do not become nonzero by parameter uncertainty of the system. This corresponds to structured uncertainty in control theory literature [25]. In general, we see that as long as the ***A*** matrix is Hurwitz and the sparsity pattern is preserved, then the system has PA. For this specific case, any positive parameters *R, C, L* makes Hurwitz ***A***. Therefore, the PA property is naturally robust to any physical parameter uncertainties in this system.

### C. RPA in control-affine systems

PA and internal model principles have also been studied in the more general setting of nonlinear input-output systems. A result relevant here is Theorem 1 in the work by E.D. Sontag [10]. It states that if system (Σ) has a uniform relative degree and satisfies certain technical conditions, then it adapting to a class of inputs implies that it contains an internal model of that class of inputs. For the case of constant disturbance, the theorem implies system (Σ) has a change of coordinates so that it can be expressed as

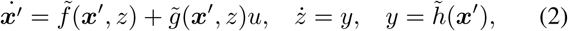

where *x′* is *n* − 1 dimensional and *z* is 1 dimensional. This implies system (Σ) satisfies IFB.

As for robustness, the system (Σ) has PA for all uncertain dynamics that keep the system stable and the integral variable *z* unchanged. For example, if *z* is only a function of *x*_1_, then any uncertain dynamics that changes *x*_2_ but keeps the system stable still satisfies PA. Compared to the efficient-to-check determinant condition for RPA in linear systems, we have to discover an integral variable and look at its particular form to determine robustness for control-affine systems.

## III. RPA IN CRN

For electrical or cyberphysical systems we can simply construct integral variables through common circuit elements or computers. In comparison, construction of integral variables is nontrivial for biomolecular reaction networks due to physical constraints on the type of dynamics chemical reactions can take.

### A. Physical constraints of CRN dynamics

We consider *n* chemical species denoted by symbols *X*_1_, …, *X_n_.* We denote their concentrations by 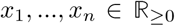. A reaction then is an expression of the form

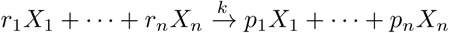

with 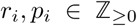 denoting the number of *X*_*i*_ molecules involved in reactants or products of this reaction. This reaction denotes reactant species on the left, product species on the right, and reaction rate 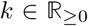 above the arrow. If we assume deterministic mass action dynamics [20], [26], then this reaction happens with propensity 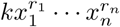, increasing *x*_*i*_ if *p*_*i*_ > *r_i_,* decreasing *x*_*i*_ if *p*_*i*_ < *r_i_,* and keeping *x*_*i*_ unchanged if *p*_*i*_ = *r_i_.* For example, reaction 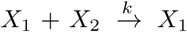 keeps *x*_*1*_ unchanged and decreases *x*_*2*_ with dynamics 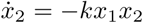. A chemical reaction network (CRN) is a finite collection of species and reactions. See [18] for more details.

Assuming deterministic mass action kinetics, the dynamics of concentration *x*_*i*_ of a chemical species have to be of the following form:

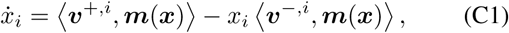

where *m(x)* is the vector of all monomials of *x*, and *v*^+^ and *v*^*−*^ are non-negative vectors with finitely many nonzero entries. 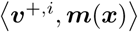 is the production propensity of *x*_*i*_, while 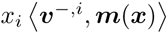 is the degradation propensity of *x*_*i*_. We see that both production and degradation have to be polynomials of *x*. More importantly, the degradation propensity of *x*_*i*_ have to be at least first order in *x*_*i*_. This is because for a well-mixed reaction system, if one molecule of *X*_*i*_ is degraded with a certain rate, then each molecule will be degraded with the same rate. So the degradation propensity of *X*_*i*_ concentration has to be at least proportional to its concentration [26], [20].

Note that this specific form is a *physical* constraint on the CRN dynamics. That means this constraint is only true for physical variables *x*_*i*_ that are concentrations of species. It is not necessarily true if we change coordinates. For example, the degradation propensity of variable *z* = *x*_1_ − *x*_2_ is a sum of the degradation propensity of *x*_*1*_ and the production propensity of *x*_2_. So it could be 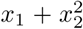, which does not have factor (*x*_*1*_ − *x*_2_).

This physical constraint is severe, in the sense that it makes it impossible to directly implement integral variables as concentrations of chemical species in many cases. This is discussed in Section III-B.

To describe the entries of *v*^*+,i*^ and *v^−,i^,* we let 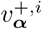 denote the entry in *v*^*+,i*^ that is coefficient of monomial 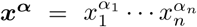 for 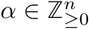 in the production polynomial, while we use 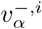 to denote the coefficient of the same monomial in the degradation polynomial. Note that (C1) then implies 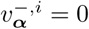 if *α*_*i*_ =0.

Another physical constraint of CRN dynamics is more subtle.

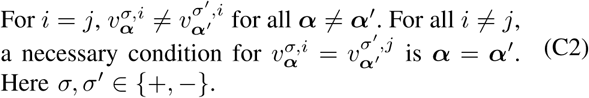

For example, we may want the production term *k*_*1*_*x*_*2*_ of *x*_*1*_ to match degradation term *k*_*2*_*x*_*2*_ of *x*_*2*_ by setting *k*_*1*_ = *k*_*2*_ so that we can cancel these two terms in dynamics of *x*_1_ + *x*_2_. This violates (C2). In contrast, any terms from the same reaction can have matching rates. For example, 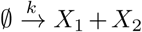 results in matching constant production terms *k* in both 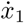 and 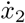, sequestration reaction 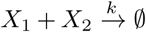 adds the term −*kx*_*1*_*x*_*2*_ in both 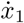 and 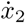, while 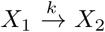 adds −*kx*_*1*_ term to 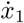 but *kx*_*1*_ term to 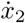.

The physical reason behind this constraint is that CRNs are mesoscopic models of biomolecular reactions [26], so the reaction rate parameters in CRNs are lumped parameters with uncertainty. Therefore, precise matching of reaction rates between different reactions is very unlikely to happen. The only way left to match terms in dynamics of two different species then is for those terms to come from the same reaction.

The significance of constraint (C2) is mostly in elimination of unreasonable solutions when we derive conditions on CRNs that satisfy PA, such as in Section III-E.

### B. Concentration adaptation

To see the effect of the above physical constraints on dynamics of CRN, for the rest of this work we consider one specific type of output: *y = x_1_ − μ* or *y = μ − x_1_,* where 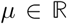 is a constant, and *x*_*1*_ is the concentration of species *X*_*1*_. This means the goal is to drive the concentration of *X*_*1*_ to a fixed constant, despite constant disturbances and uncertain dynamics. We call this concentration adaptation. For the rest of the paper, we will take *y = μ − x_1_,* as almost all results carries over to the case *y = x_1_ − μ.* We make the additional assumption that *x*_*1*_ is not an integral variable itself to make the problem nontrivial.

Concentration adaptation has been the central focus of studies on perfect adaptation in biomolecular systems [23], [12], [6]. Other topics of interest can often be formulated as a close variant of a concentration adaptation problem. For example, the fold change detection problem can be formulated with output *y* = μ − log *x*_1_, where log is approximately implemented by allosteric proteins [27]. Similarly, concentration tracking can be formulated with output *y* = *w* − *x*_1_, so *x*_1_ tracks input *w*.

With the concentration adaptation control objective in mind, we immediately see that integral variables cannot be implemented via a chemical species. This is because both 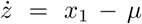 and 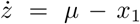 do not satisfy the physical constraint (C1) if *z* is concentration of a chemical species other than *x*_1_. This forces integral variables to be implemented in CRN network architectures.

### C. Implementation of integral variables in CRN

Here we describe three general ways of implementing integral variables in CRN and illustrate each through network motifs well-known in systems biology. Network motifs are diagrammatic representations of circuits widely used as guiding principles for controller design in systems biology and synthetic biology [17], [23].

1) *Constrained integral variable:* One way to approximately implement an integral variable is to relax the constraint 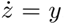. We want the concentration of some chemical species satisfy 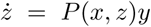, so *z* reaching steady state does not in general guarantee *y* = 0 everywhere. This makes integral variables directly implementable by chemical species, but sacrifices the effectiveness of integral variables achieving the control objective.

We consider an example from cell population control that also uses this approach [4], [2], [3]. The CRN is the following:

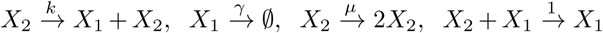

Here *X*_*2*_ is the number of cells that self-replicates, and *X*_*1*_ is a toxin that kills the cells. The dynamics satisfy

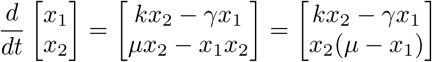

We see that *x2* acts as the integral variable for *y* = μ − *x*_*1*_ with additional steady state *x_2_ =* 0.

2) *Change of coordinates:* Another implementation method relaxes the constraint that *z* is a chemical species. We take *z* = *ℓ(x),* so *z* is a virtual variable. One network motif that is an example of this approach is incoherent feedforward (IFF) [23]. The simplest such example is the “Sniffer” model [15] with the following reactions:

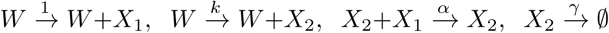

Here *W* acts as an enzyme that produces proteins *X*_*1*_ and *X*_2_, while *X*_*2*_ is in turn a protease that degrades *X*_1_. This system has dynamics

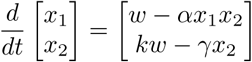

At steady state, 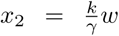, and 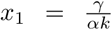. So *x*_*1*_ is independent of *w* and initial conditions. This system has PA for concentration tracking output *y* = μ − *x*_*1*_ where 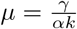. Consider *z = kx_1_ − x_2_,* we have 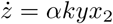. So *z* acts as a constrained integral variable.

*3) Approximation:* The third approach is to circumvent the physical constraint (C1) by approximations, including time-scale separation and asymptotics in large parameters. One network motif that is an example of this approach is called integral negative feedback (IFB) [12], [28].

Consider the following reaction network:

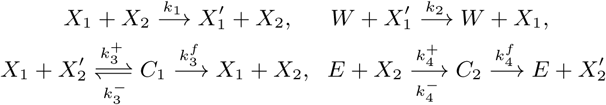

*X*_*i*_ and 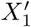 are two states of the same protein, *X*_*2*_ and 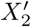 are two states of another protein, *C*_*1*_ and *C*_2_ are intermediate complexes, *W* is input, an enzyme that catalyzes the transformation of 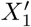 to *X*_*1*_, and *E* is some external enzyme that catalyzes the transformation of *X*_*2*_ to 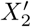.

This system has the following dynamics after taking quasi steady state approximations to obtain Hill functions [1]:

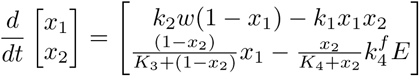

where 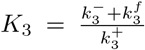, and 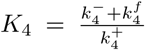. In addition, we normalized units of *x*_*i*_ and *x*_*2*_ to make 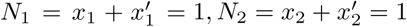, and normalized time so that 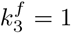.

In the limit that *K*_3_, *K*_4_ are very small, we have approximation 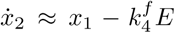. So, at steady state, 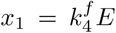, independent of *w.* Here *x*_*2*_ acts as the integral feedback variable for output 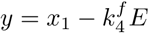.

Note that since *K*_*3*_ and *K*_*4*_ are assumed to be small compared to *x*_*2*_ and *1 − x_1_*, we need the steady state of *x*_*2*_ to be far from 0 or 1. While *x*_*1*_ steady state does not depend on *w, x*_*2*_ steady state does (it is 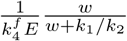). This puts a constraint on what the constant disturbances can be.

### D. Connection with Absolute Concentration Robustness

In addition to systems biology and synthetic biology literature, a special class of PA called absolute concentration robustness (ACR) has also been studied using very different approaches in the field of chemical reaction network theory (CRNT) [18], [19]. ACR is relevant for CRNs with conservation classes. For example, in *X*_*1*_ ⇌ *X*_*2*_, *x*_1_ + *x*_2_ = *N* is conserved during time evolution of the system. ACR then describes the property that the concentration of one chemical species has a positive steady state that is independent of the conserved quantity *N*. Through an example, we show how our constraint-focused perspective with general implementation methods can be used to understand ACR as well.

Consider the following CRN that satisfies ACR:

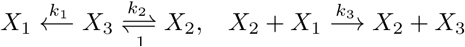

Note that there is a conserved quantity *x_1_ + x_2_ + x_3_ = N* as no molecules are created or destroyed in this CRN. ACR theory then states that species *X*_*1*_ has ACR, meaning *x*_*1*_ has a positive steady state that is independent of the total concentration *N*. The dynamics of the system satisfy

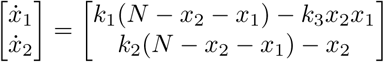

Let 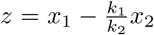, then *z* has dynamics 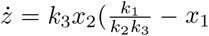.

So *z* acts as a constrained integral variable for output *y* = *μ* − *x*_*1*_ where 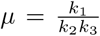. This system uses both the linear change of coordinates method and the constrained integral variable method. Note that ACR is a special case of PA, as PA could reject disturbances other than the total concentration.

**Fig. 2.**
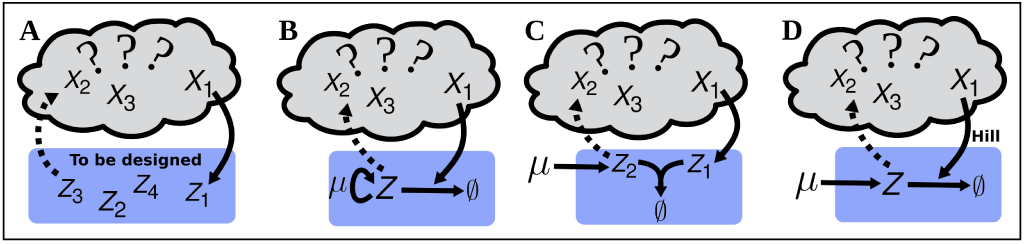
Graphical depiction of various constructions of integral variable for unknown plants with the concentration adaptation output. (A) Graph showing the general task of designing an integral variable for an unknown plant, assuming we can have reactions involving *X*_*1*_ (solid arrow), while postponing feedback actuation for future considerations (dashed arrow). (B) Integral variable constructed via the constrained integral method. Here *Z* self-replicate itself with rate *μ,* and *X*_*1*_ catalyze degradation of *Z* with rate 1. (C) Integral variable constructed via the change of coordinate method. Here *Z*_1_ and *Z*_2_ annihilate each other, *Z*_2_ is produced at constant rate *μ,* and *X*_*1*_ catalyze production of *Z*_*2*_ with rate 1. (D) Integral variable constructed via the approximation method. Here *Z* is produced at constant rate *μ,* and *X*_*1*_ acts as an enzyme that catalyzes degradation of *Z* through a first order Hill function kinetics.

### E. Integral variable design for unknown plants

Previous studies in biological control mostly focused on discovering integral variables in specific circuits found in nature. Because there is no clear divide between the plant and the controller in a closed loop system, the constructions of integral variables previously found usually depend heavily on the specific biomolecular mechanisms of that system [12]. This dependence has limited the use of these constructions in biological control.

Following the separation of controllers and plants in control theory, we consider the problem of constructing integral variables for arbitrary plants, which is useful for biological control. This is easy in electrical and cyberphysical systems, but nontrivial and relatively unstudied in biological control. Indeed, the first success in finding an integral variable construction for arbitrary plants in 2016, the sequestration feedback controller, has led to theoretical and experimental progress in biological control [6].

To illustrate the power of the “forward” approach taken in this paper, we show how we can analytically solve for such constructions. We derive two new constructions of integral variables for unknown plants that has biological relevance.

The general situation we consider is illustrated in Figure 2-A. For unknown plants, we want to design an integral variable implemented by controller species *Z*_*1*_,…,*Z*_*m*_ to drive one plant species *X*_*1*_ to a fixed concentration *μ* that we can tune externally. This corresponds to designing CRN dynamics of chemical species *Z_1_,…,Z_m_.* Since we only know that the plant will contain species *X*_*1*_ while knowing nothing else about the plant, dynamics of *z*_*i*_’s can only depend on each other and *x*_1_, but nothing else. From (C1), we have

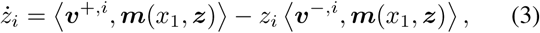

where *i* = 1,…, *m*, 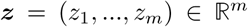 is the vector of concentrations of species *Z_1_,…, Z_m_,* and *m(x*_1_, *z)* is the vector of all monomials in *x_1_,z_1_,…, z_m_.*

The goal is to use change of coordinates, constrained integral feedback, and approximation methods to find *z*\* = *ℓ(z_1_,…, z_n_)* for some function *ℓ* such that 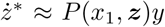 for some polynomial *P*, where *y* = *μ − x*_1_. In general, the solution set is too large to be useful for biological construction, so we want to pose additional constraints to focus on solutions that are implementable and interpretable. In the following, we focus on deriving the simplest possible integral variable constructions in CRN using only one of the three approaches.

1) *Constrained integral feedback:* The easiest to derive is the constrained integral variable method. Since we do not use approximations or change of coordinates, we require *z*\* to be the concentration of one of the *Z*_*i*_ chemical species with dynamics 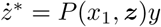.

We consider *m =* 1, so we only have one controller chemical species, *Z*. Its concentration, *z*, has dynamics

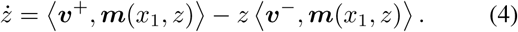

It can be shown through routine calculations that 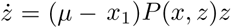 satisfies the form of (4) for any polynomial *P*, while no lower degree production or degradation terms does. Therefore, taking *P(x, z)* to be 0 degree, i.e. a constant, gives the simplest possible implementation. Focusing on the *y* = *μ* − *x*_*1*_ case, we have z dynamics

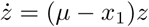

which can be implemented by the simple chemical reactions

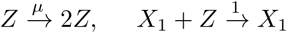

A graphical depiction of this integral variable design is shown in Figure 2-B. Note that population control circuits are often special cases of this design [4], [3], [2].

2) *Change of variable:* The sequestration feedback construction of an integral variable uses change of coordinates [6]. We can derive this construction by restricting to linear change of coordinates and posing constraints on the number of controller species allowed (*m*) as well as on the degrees of monomials allowed for production (*d*_*p*_) and degradation (*d*_*d*_) terms. Since we do not use approximations or constrained integral variables, the goal is to find 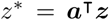 such that 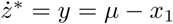.

We know that *m* = 1 is infeasible, as *z*\* cannot be *x*_1_. So the lowest possible *m* is 2. For this case, we can take *z*\* = *z*_*1*_ + *az*_*2*_ for constant 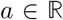, then according to (C1), *z*^*\**^ satisfy

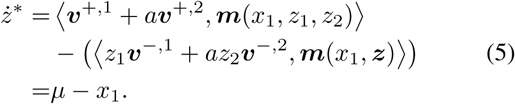

In terms of notation, we use 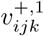 to denote the coefficient of monomial 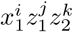 in the production polynomial of *z*_1_, and 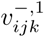 to denote the coefficient of the same monomial in the degradation polynomial of *z*_2_. Recall that (C1) forces 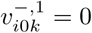, for example.

With (5), we can look at the polynomials on the two sides of the equation and match the coefficients term by term. For *d_p_ =* 0, there can be no linear term in *x*_1_, so the lowest possible *d*_*p*_ is 1. Taking *d*_*p*_ = 1, *d*_*d*_ = 0, we find that (5) can only be satisfied if there is no degradation at all (e.g. 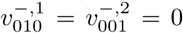), or if there is matching parameters that cannot be from the same reaction, i.e. 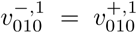 and 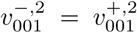. The former results in instability, while the latter violates constraint (C2).

So we are forced to look at *d*_*p*_ = *d*_*d*_ = 1. For this case, the only way for (5) to be satisfied without instability or violation of (C2) is to have 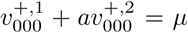, 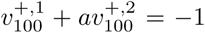, and the other parameters zero. Among these, 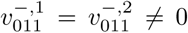 is allowed if they come from the same reaction *Z*_1_ + *Z*_2_ → *C*, where *C* is some set of species not including *Z*_1_ and *Z*_2_, such as 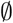 or *X*_1_.

The following set of reactions satisfies the above:

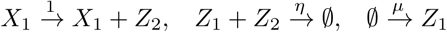

Here *X*_1_ catalyzes the production of *Z*_2_, *Z*_1_ is produced with a constant rate *μ* that is our reference, and *Z*_1_ together with *Z*_2_ annihilate each other. This design is found by ingenuity in [6], while we can formally derive it. A graphical depiction of this design is shown in Figure 2-C.

*3) Approximation:* The goal here is to find *z*\* that is the concentration of one of the *Z*_*i*_ chemical species with dynamics 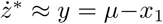, since we use only approximation.

To systematically search for possible constructions of integral variables, we need to search through dynamics that are possible through time scale separation and large parameter asymptotics. This can be done with the help of general theories on time scale separation in CRNs developed in, e.g., [29], [30], and is a goal for furture work. Here we restrict ourselves to first degree Hill functions that are readily implementable by enzymatic reactions that satisfy Michaelis Menten dynamics.

We again consider the simplest possible case where *m* = 1, so *z*\* = *z*, concentration of the only controller species *Z*. In addition, we require *d*_*d*_ = *d*_*p*_ = 0, with Hill function degradation term catalyzed by *x*_1_. This gives

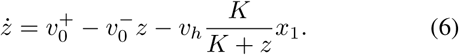

By taking 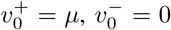, *v*_*h*_ = 1, and the approximation that *K* is very small compared to *z*, we have 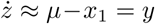. The dynamics of *z* can be implemented by the following reactions:

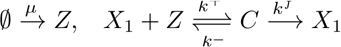

Here *Z* is produced with constant rate *μ*, and *X*_1_ acts as an enzyme that catalyzes the degradation of *Z*. Under quasisteady state approximation [1], *z* satisfy (6) with *v*_*h*_ = *k*^*f*^ and 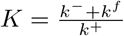. The graphical depiction of this construction is shown in Figure 2-D.

The above examples show that we can systematically derive constructions of integral variables for unknown plants, starting from constraint (C1) with the help of constraint (C2). When we know more about the plant, such as when we know *X*_*2*_ produces *X_1_,* we can perform similar calculations to derive constructions of integral variables suitable for these cases.

### F. Integral variable design for incoherent feedforward

The drawback of the above approach in integral variable design is that it always gives rise to a feedback system. This is because the integral variable is constructed by controller species outside the plant, as the plant is unknown or unmodifiable. Hence the integral variable has to be fed back to the plant to be effective. Technically, feedback is usually required to guarantee that the closed loop system is stable. However, many regulatory networks with PA found in biological systems are of the form called incoherent feedforward (IFF), which does not have feedback [28], [23]. IFF networks have the input disturbance influencing two branches of the network that both influence the target species, one activating, and one inhibiting. IFF networks then need to guarantee that the magnitude of the activating branch matches that of the inhibiting branch through structured physical interconnections, so as to obtain perfect adaptation.

Due to this canceling mechanism of IFF, it can only be obtained by a design problem where we are given how the disturbance enters the target variable while we can design the entire closed loop system. Since we need to construct a feedforward system that has a feedback interpretation, we have to use the change of coordinates method as it is the only one of the three methods that can transform a feedforward system into a feedback one. Furthermore, to design an IFF system we need to design a closed loop system with stability considered, which is distinctively different from the integral variable design problem with unknown plants where design of feedback actuation and stability are postponed to the moment when we know more about the plant.

We show below that we can derive IFF structures through similar procedures as in integral variable design for unknown plants in Section III-E.

Consider the simplest IFF design problem where disturbance *W* enters plant species *X*_*1*_ by catalyzed production 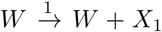. Since we do not modify how disturbance enters *x_1_,* by (C1) *x*_*1*_ has dynamics

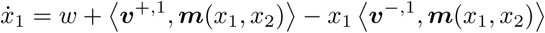

The simplest inhibiting branch would have one additional plant species, call it *x*_2_, with dynamics

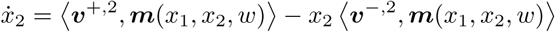

Note that the monomials include *w* in dynamics of *x*_2_ as we can design how disturbance enters in the inhibiting branch.

Assuming we use linear change of coordinates and constrained integral variables but not approximations, the goal is to find *z* = *x*_*1*_ + *ax*_*2*_ with constant 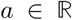 such that 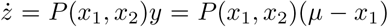 for some polynomial *P*.

Now we can again constrain the degree of monomials in the production *(d_p_)* and degradation (*d*_*d*_) terms to find the simplest possible design. The (*d_p_,d_d_*) = (0,0) and (1,0) cases are impossible as both require *x*_2_ to have a vanishing degradation propensity. The (0, 1) case is impossible as disturbance *w* cannot enter *x*_2_ to construct *z*. So the simplest case is the (*d_p_,d_d_*) = (1,1) case, where the system dynamics satisfy

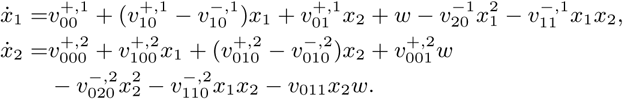

Looking at 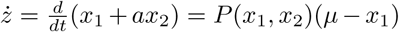, we can just match coefficients term by term to see the constraints. If *P(x_1_,x_2_) = P(x_1_*), the equation for 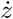 cannot be satisfied as it again requires all degradation terms of *x*_*2*_ to have coefficient 0 if we satisfy constraint (C2). So the simplest case beyond these is *P(x_1_,x_2_) = x_2_.* For this case the constraints turns out to be 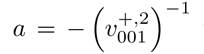 to cancel out the *w* term, 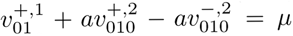 for the *x*_*2*_ term, and 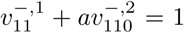 for the *x*_*1*_*x*_*2*_ term. One special case that satisfies the above is the “Sniffer” model presented in Section III-E.2.

We see from this example that we can systematically derive CRNs that satisfy IFF as well, given how the disturbance enters one branch of the network.

## IV. DISCUSSION

In this work we investigated a control theory framework to understand perfect adaptation in biomolecular reaction networks. This framework is of a “ground-up” or “forward engineering” flavor, with strong focus on the physical constraints on the dynamics of chemical reaction networks. Considering the task of concentration adaptation, we showed that physical constraints force biological systems to encode perfect adaptation properties in the network architecture instead of dynamics of some chemical species. We then developed systematic ways to derive reaction networks with perfect adaptation and constructions of integral variables for unknown plants.

We saw that the severe constraints on possible dynamics enabled rather different approaches to control problems for biomolecular systems. Notably, constraints on possible controller dynamics is highly relevant not only to biological control. For example, in legged robotics, hard constraints built into the robot by good mechanical design could result in much easier to control robots (see page 25 in [31]). However, in contrast with cyberphysical controller design where all calculations are done in computers, the design of these hard constraints falls into this category of controller synthesis with severe physical constraints. For control problems where the cyberphysical approach of sensor-computer-actuator work flow encounters difficulties, controller synthesis that results in physically implementable dynamics may point to alternative solutions.

In this work, we mostly focused on methods that can derive integral variables that are implementable by chemical reactions. It should be emphasized that designing an integral variable is only the first step of solving the concentration adaptation problem. The system only performs as desired if we also have stability, which is mostly ignored in integral variable design. In general, we also need to design feedback actuations that use integral variables to actuate the plant to guarantee stability (the dashed arrows in Figure 2). These feedback actuation designs are rooted in characterizations of stability, which is an important topic to be studied further with this physical constraints based framework.

As was discussed in Section II, the PA property of a CRN is naturally robust to uncertainties that preserve stability of the system and structure of the integral variable. This has nontrivial biological implications. For example, for the population control CRN described in Section III-E.1, the detailed dynamics of the toxin does not matter as long as the cell population dynamics is intact and the system is stable.

Lastly, the physical constraints this work focused on can act as a parameterization scheme of possible controllers with perfect adaptation property. This parameterization together with nonlinear controller synthesis methods [32] may lead to a general controller synthesis method for chemical reaction systems.

